# Repurposing nystatin and taurocholate to target intracellular spore persistence during antibiotic treatment to prevent recurrent *Clostridioides difficile* infection

**DOI:** 10.64898/2026.09.25.754266

**Authors:** Pablo Castro-Córdova, María Mendoza-León, Said Álamos-Musre, Nicolás Montes-Bravo, Juan Muñoz-Miralles, Fernando Díaz-Yáñez, Felipe M Llancalahuén, Diego Cea, Rosario Hernández-Armengol, Javiera Valeria, Katherine Mondaca, Carolina Otero, Fernando Gil, Marjorie Pizarro-Guajardo, Daniel Paredes-Sabja

**Affiliations:** ANID – Millennium Science Initiative Program – Millennium Nucleus in the Biology of the Intestinal Microbiota, Santiago, Chile; IMPACT, Center of Interventional Medicine for Precision and Advanced Cellular Therapy, Santiago, Chile; Departamento de Medicina Interna, Faculty of Medicine, Universidad de los Andes, Santiago, Chile; Programa de Oncología Traslacional, Centro de Investigación e Innovación Biomédica (CiiB), Universidad de los Andes, Santiago, Chile; Escuela de Química y Farmacia, Facultad de Medicina, Universidad Andrés Bello, Santiago, Chile; Microbiota-Host Interactions & Clostridia Research Group, Center for Biomedical Research and Innovation (CIIB), Universidad de los Andes, Santiago 7620001, Chile; School of Medicine, Faculty of Medicine, Universidad de los Andes, Santiago 7620001, Chile; Department of Biology, Texas A&M University, College Station, Texas, USA

**Author notes:** Corresponding Author. Dr. Daniel Paredes-Sabja, Department of Biology, Texas A&M University, College Station, TX, 77843, USA.

**Keywords:** Nystatin, *C. difficile* spores, taurocholate, Recurrent CDI, spore persistence, *C*. *difficile* treatment

## Abstract

Successful treatment of primary *Clostridioides difficile* infection (CDI) while preventing its recurrence remains a major clinical challenge, with recurrent CDI (R-CDI) affecting 20–30% of treated patients. Persistence of *C. difficile* spores within the intestinal mucosa is thought to drive R-CDI following antibiotic treatment. Here, we show that intracellular *C. difficile* spores remain protected from taurocholate-triggered germination, and that nystatin, an FDA-approved antifungal that blocks spore entry into host cells, reduces this protected intracellular population and redirects spores toward an extracellular, germinating population accessible to antibiotics. In a murine model of R-CDI, oral co-administration of taurocholate with nystatin and antibiotics during primary CDI reduced R-CDI incidence from 100% to 44%. When administered during an active R-CDI episode, the same formulation reduced a second R-CDI from 100% to 62%. To improve bioavailability, we encapsulated nystatin and vancomycin in chitosan nanoparticles. Oral administration of nanoparticle-encapsulated vancomycin, alone or combined with nanoparticle-encapsulated nystatin, reduced R-CDI colonization to 33% and 44%, respectively, compared with 100% in mice receiving free vancomycin. Together, these findings support a therapeutic strategy that repurposes nystatin and taurocholate, in combination with antibiotics and nanoparticle-based delivery, to target intracellular spore persistence and reduce the incidence of *C. difficile* recurrence..

**Graphical abstract:** Combination of Vancomycin, nystatin and taurocholate protects from recurrence of CDI

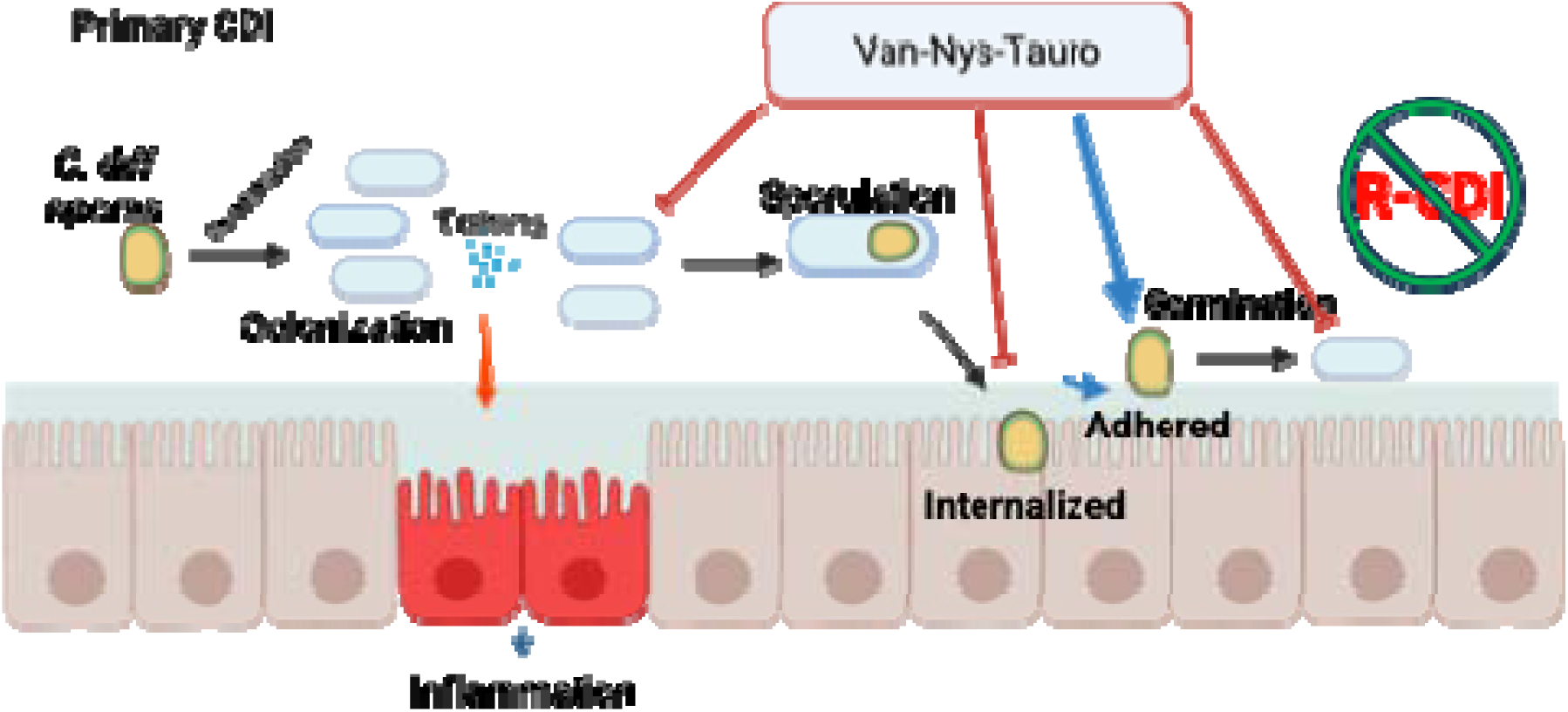

## INTRODUCTION

*Clostridioides difficile* is the most common cause of nosocomial diarrhea and a life-threatening infection ^1–3^. The Centers for Disease Control and Prevention (CDC) has designated *C. difficile* an urgent health threat despite updated infection control practices ^4^. Even after appropriate therapy, ∼20–35% of patients develop recurrent CDI (R-CDI), reappearance of symptoms within 2–8 weeks after treatment completion, with further recurrences spiking to 65% ^1, 3, 5–7^. Mortality rates increase from ∼8% during primary CDI to ∼20–30% with R-CDI ^8–10^, underscoring the need for novel therapeutic approaches ^11^.

*C. difficile* spores germinate in patients with a dysbiotic microbiota during infection ^12, 13^, while colonization produces toxins A and B (TcdA, TcdB), responsible for clinical symptoms ^14, 15^. Simultaneously, *C. difficile* forms highly resistant dormant spores essential for transmission and recurrence ^16–18^. Vancomycin and fidaxomicin, the current standard of care, resolve most primary CDI cases ^19, 20^, with fidaxomicin yielding nearly half the R-CDI rate of vancomycin ^19, 21, 22^. However, recovered patients continue to shed fecal spores ^18^, underscoring the need for improved spore-targeting.

Several strategies have entered clinical trials to address *C. difficile* persistence. SER-109 (VOWST), a purified Firmicutes spore consortium, reduced R-CDI to 12% versus 40% with placebo in phase 3 trials ^23^. VP20621 (NTCD-M3), a nontoxigenic *C. difficile* strain providing competitive colonization resistance, reduced recurrence to 11% versus 30% with placebo in a phase II trial ^24, 25^. Ramoplanin, which inhibits cell wall synthesis by blocking Lipid I-to-Lipid II conversion and peptidoglycan transglycosylation ^26, 27^, binds *C. difficile* spores persistently and inactivates them after taurocholate-triggered germination ^28^. Passive oral spore-specific immunotherapy has also reduced R-CDI in preclinical models ^29^. Aside from SER-109, available only by prescription at $19,000–21,000 per 3-day course ^30^, none of these approaches has achieved regulatory approval or clinical adoption, underscoring the need for effective, accessible R-CDI prevention.

Bezlotoxumab (Merck), an FDA-approved monoclonal antibody targeting TcdB, reduced R-CDI rates from 27% to 17% when given intravenously alongside standard antibiotic therapy ^6, 31^, but was discontinued in January 2025 after failing a phase 4 trial ^32^. The unmet need for accessible R-CDI therapeutics remains urgent.

We previously showed that *C. difficile* spores internalize into the intestinal mucosa, potentially forming a dormant intracellular reservoir that persists to cause R-CDI ^33^. Nystatin, an FDA-approved antifungal safe in humans ^34^, inhibited spore entry into intestinal epithelial cells *in vitro* and delayed R-CDI onset by ∼20% when combined with vancomycin ^33^, suggesting that blocking spore entry during CDI could prevent this reservoir and delay recurrence.

We hypothesized that preventing epithelial spore internalization while forcing extracellular germination would expose the protected spore reservoir to antibiotic killing, reducing R-CDI. Using fluorescence microscopy, we show that nystatin blocks *C. difficile* spore internalization into intestinal epithelial cells, enabling subsequent taurocholate-triggered germination. In a murine R-CDI model, co-administration of taurocholate, nystatin, and vancomycin reduced subsequent recurrence. Encapsulating nystatin and vancomycin in chitosan nanoparticles further improved this formulation, reducing recurrence after a primary or first recurrence by 33% and 44%, respectively. These results introduce a novel therapeutic strategy, targeting intracellular *C. difficile* spores by repurposing nystatin and taurocholate alongside vancomycin or other antibiotics, and provide a framework for developing safe, standardized treatments targeting spore persistence in the intestinal mucosa.

## METHODS

### *Clostridium difficile* Strains and Spores Purification

The *C. difficile* R20291 CM210 strain (derived from R20291, Nigel Minton) was cultured anaerobically at 37°C in BHIS broth or agar. For sporulation, cultures were plated on 70:30 sporulating agar and incubated 7 days at 37°C; colonies were harvested in ice-cold Milli-Q (MQ) water and pelleted by centrifugation. Spores were purified by density-gradient centrifugation over 45% Nycodenz, as previously described ^35^, washed repeatedly in MQ water, enumerated, adjusted to 5×10 spores/mL, and stored at −80°C (extended methods in *Supplemental Information*).

### Cell culture

Caco-2 cells (ATCC) were cultured at 37°C with 5% CO in DMEM high glucose supplemented with FBS, penicillin, and streptomycin, and seeded onto glass coverslips 2 days pre-confluence for infection experiments (extended methods in *Supplemental Information*).

### Germination assay of *C. difficile* spores pure and in Caco-2 monolayers

Germination of heat-activated (60–70°C) pure or Caco-2-associated spores was assessed by phase-contrast microscopy after incubation with taurocholate and glycine. To distinguish internalized from extracellular spores, infected monolayers were double-stained before and after permeabilization: extracellular spores were labeled with anti-*C. difficile* spore goat serum plus donkey anti-goat CF568 in non-permeabilized cells, while total spores (extracellular plus internalized) were labeled with the same primary antibody plus donkey anti-goat CruzFluor488 after permeabilization (PBS-0.2% Triton X-100); internalized spores were defined as total-positive but extracellular-negative. To test whether nystatin reduces the internalized, germination-protected population, monolayers were pretreated with 30 µM nystatin before infection with NHS-opsonized spores and taurocholate/glycine challenge. Samples were imaged on an Olympus BX53 fluorescence microscope (100× oil objective) with a Qimaging Retiga R6 camera, and spores were counted using the Cell Counter plugin in ImageJ (extended methods in *Supplemental Information*).

### Preparation of chitosan nanoparticles

VAN-CS-NPs and NYS-CS-NPs were synthesized by ionic gelation. A 1% low-molecular-weight chitosan solution (75–85% deacetylated) in 1% acetic acid was combined with vancomycin (0.755 mg/mL) or nystatin (0.0755 mg/mL) and added via syringe pump into 1.55 mg/mL pentasodium tripolyphosphate (TPP) under stirring; nanoparticle formation was indicated by a transparent-to-opaque transition. Nanoparticles were sonicated, centrifuged, and resuspended in ultrapure water. Size, polydispersity index (PDI), and zeta potential were determined by dynamic light scattering (Nano-ZS Zetasizer, Malvern) across three independent batches per formulation (extended methods in *Supplemental Information)*.

### Lyophilization of nanoparticles

VAN-CS-NPs and NYS-CS-NPs were lyophilized with 40 mg/mL sucrose and stored at −20°C until use (extended methods in Supplemental Information).

### Encapsulation efficiency

Encapsulation efficiency was determined by HPLC quantification of vancomycin or nystatin remaining in nanoprecipitation supernatants. Based on these measurements, we produced 8 batches of VAN-CS-NPs (30.93 mg final mass) and 16 batches of NYS-CS-NPs (73.44 mg final mass) for *in vivo* use (extended methods in Supplemental Information).

### Mice

Female and male C57BL/6 mice (8–12 weeks old) were housed individually under conventional conditions with *ad libitum* food and water. All protocols were approved by the Institutional Animal Ethics Committee of Universidad Andrés Bello (#022/2016, #007/2019, #018/2019) and by the Institutional Animal Care and Use Committee (IACUC) of Texas A&M University (extended methods in Supplemental Information).

### R-CDI mouse model

Mice received an antibiotic cocktail (kanamycin, gentamicin, colistin, metronidazole, vancomycin) by gavage for 3 days, followed by intraperitoneal clindamycin, then infection with *C. difficile* spores or vegetative cells, as previously described ^33, 36, 37^. Investigators were not blinded, to avoid cross-contamination between treatment groups. Mice were monitored daily for weight, appearance, behavior, and diarrhea (scored 0–3, with a score ≥1 considered diarrhea) and euthanized at experimental endpoint for collection of cecal content and intestinal tissue ^33, 38^.

For the vancomycin/ramoplanin/nystatin experiment, mice (6×10 spores) were treated from days 3–8 post-infection with i) vancomycin (50 mg/kg; *n*=10), ii) vancomycin plus nystatin (17,000 IU/kg; *n*=13), or iii) vancomycin plus ramoplanin (8 mg/kg) plus nystatin (*n*=15). For the taurocholate-formulation experiment, mice received i) vancomycin (*n*=14) or ii) vancomycin plus ramoplanin plus nystatin plus taurocholate (20 mg/kg; *n*=11) from days 3–8. To model a second R-CDI, mice (*n*=30) were treated with vancomycin from days 3–8; those developing CDI and R-CDI (*n*=15) were then treated from days 13–17 with i) vancomycin (*n*=7) or ii) the four-drug formulation (*n*=8). For the nanoparticle experiment, mice (*n*=34) were treated from days 3–8 with i) vancomycin (*n*=9), ii) VAN-CS-NPs (equivalent to 50 mg/kg vancomycin; *n*=6), iii) vancomycin plus nystatin (*n*=9), or iv) VAN-CS-NPs plus NYS-CS-NPs (equivalent to 17,000 IU/kg nystatin; *n*=9) *(extended methods in Supplemental Information)*.

### Quantification of CFU of *C. difficile* from feces and intestinal tissues

Fecal samples and intestinal tissues (proximal colon, middle colon, distal colon, cecum) were ethanol-treated, serially diluted, and plated on TCCFA agar under anaerobic conditions; colonies were expressed as Log (CFU/g). Time to colonization was defined as the day fecal spore counts reached ≥1×10 CFU/g during the second R-CDI (*extended methods in Supplemental Information)*.

### Cecum content cytotoxicity assay in Vero cells of infected mice

Cytotoxicity was assessed as previously described ^33^. Cecal contents were clarified, filtered, serially diluted, and applied to confluent Vero cell monolayers; cytotoxic titer was defined as the reciprocal of the highest dilution producing rounding in ≥80% of cells, adjusted per gram of cecal content (*extended methods in Supplemental Information*).

### Statistical Analysis

Prism 9 (GraphPad Software, Inc.) was used for statistical analysis. Relative weight, diarrhea score, and CFU/g of feces were evaluated using the Kruskal-Wallis test with Dunn’s post hoc test; time to diarrhea and time to colonization were evaluated using the Log-rank (Mantel-Cox) test; CFU/g of tissue and cytotoxicity of cecal content were assessed using a two-tailed Mann-Whitney test.

## RESULTS

### Nystatin inhibition of spore internalization allows protected spores to germinate with taurocholate

During vancomycin treatment of *C. difficile*-infected mice, fecal taurocholate and cholate concentrations increase ^39^, facilitating spore germination in the gut ^40, 41^. We previously showed that nystatin, an inhibitor of spore entry into host cells, increases taurocholate-induced germination ^33^, suggesting that internalized spores are protected from taurocholate-triggered germination. We therefore hypothesized that internalized spores would remain phase-bright (dormant) despite extracellular taurocholate and glycine. Since heat-activation sensitizes spores to germinate with taurocholate [38], we tested whether heat-activated spores (60 °C, 30 min) germinated after infecting Caco-2 monolayers for 1, 2, or 3 h, followed by incubation with 5 mM taurocholate and 1.3 mM glycine (Fig. 1A, Fig. S1). Phase-contrast microscopy showed that after 3 h, ∼27–23% of spores in Caco-2 cells remained phase-bright versus ∼7–9% in PBS (Fig. 1B), indicating that a subpopulation of spores remains protected from taurocholate-triggered germination inside intestinal epithelial cells.

**Figure 1.**
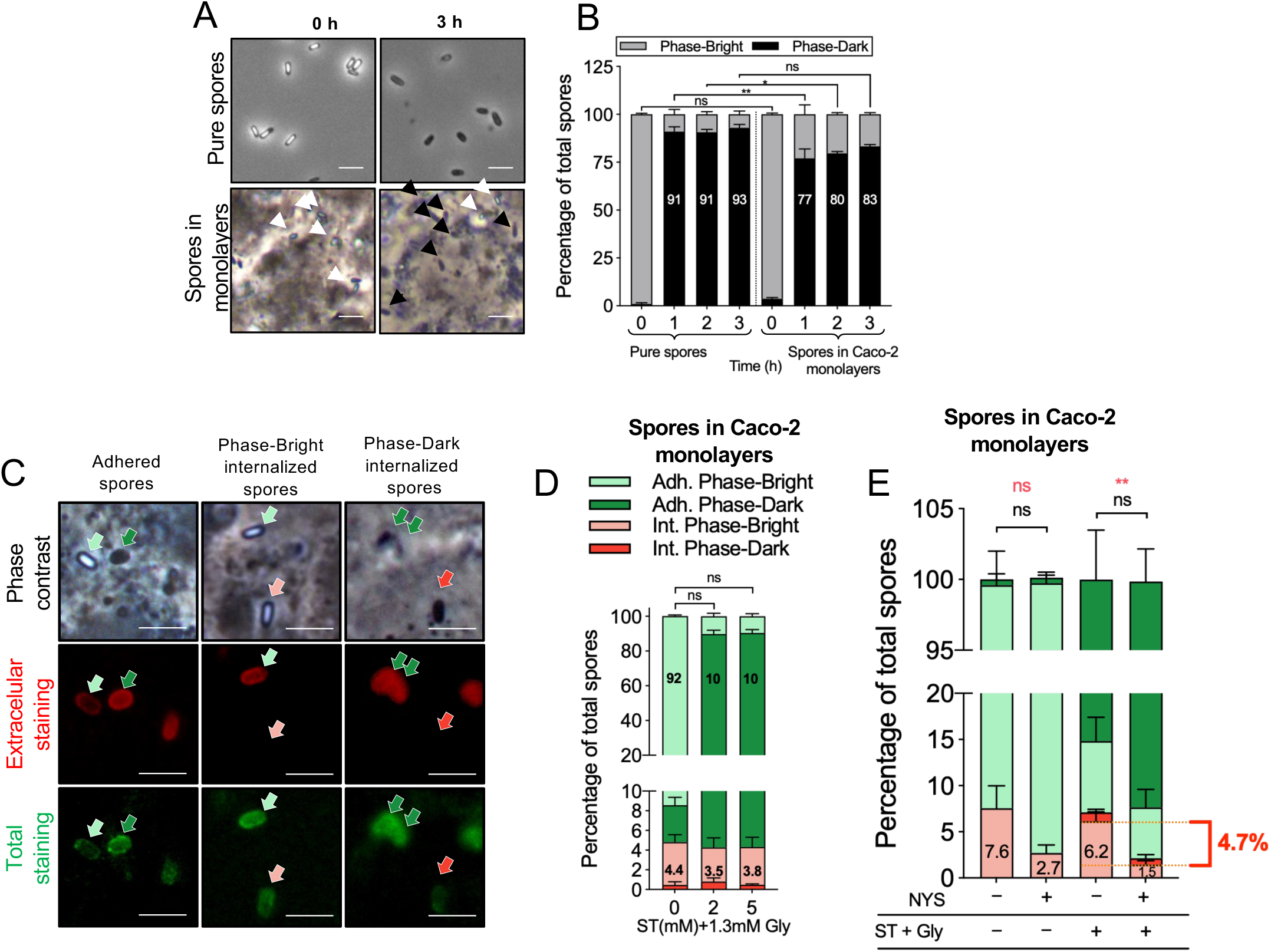

To determine whether epithelial-protected spores remained phase-bright with taurocholate, we infected Caco-2 cells with spores, added taurocholate (2 or 5 mM), and used double-staining to distinguish internalized from extracellular spores: adhered spores were tagged with anti-spore antibody and red fluorophore before permeabilization, and total spores with anti-spore antibody and green fluorophore after permeabilization (Fig. 1C). Micrographs showed that phase-bright (dormant) and phase-dark (germinated) spores were both present in epithelial cells; ∼95% of spores were adhered to the cell surface independently of germinant, indicating no role for taurocholate in internalization. Adhered germinated (phase-dark) spores reached ∼2% without germinant but ∼85% with either taurocholate concentration (Fig. 1D), leaving ∼10% of adhered spores dormant, suggesting most adhered spores remain susceptible to germination. Internalized spores comprised 4% of total spores, of which >95% remained phase-bright regardless of germinant (Fig. 1D), showing that intracellular spores stay isolated from taurocholate.

Since nystatin, a cholesterol-sequestering agent, inhibits spore internalization, we tested whether blocking internalization reduces the fraction of dormant spores. Nystatin pretreatment reduced internalized spores from ∼6–7% to ∼2–3% (Fig. 1E); combining nystatin with taurocholate reduced intracellular-dormant spores from 6% to 1.5% (Fig. 1E). Among extracellular adherent spores, addition of germinant triggered germination in almost 90% of spores; nystatin reduced the dormant fraction to 8% (Fig. 1G). These results indicate that nystatin pretreatment followed by germinant addition reduces the intracellular dormant-spore reservoir.

### Vancomycin treatment in combination with ramoplanin and nystatin delays onset of diarrhea in R-CDI mice model

*C. difficile* spores can persist viable in the gastrointestinal tract despite antibiotic treatment ^5, 18, 43^ and drive infection recurrence ^17^. Combined with our finding that internalized spores resist taurocholate-triggered germination, this suggests that surviving spores may be released after treatment ends and recolonize the host. We previously showed that oral nystatin combined with vancomycin delays recurrence in a murine model ^33^. Ramoplanin, a glycolipodepsipeptide antibiotic active against vegetative C. difficile, binds the spore exosporium for a prolonged period, positioning it to attack cells as they germinate ^28^, suggesting its addition could target persisting spores and help reduce recurrence. Antibiotic-treated mice were infected with *C. difficile* spores (Fig. 2A); all three groups showed similar weight loss, diarrhea onset, and spore shedding during CDI (days 1–3; Fig. S2A–C). From day 3, mice were treated for 5 days with i) vancomycin, ii) vancomycin plus ramoplanin, or iii) vancomycin, ramoplanin, and nystatin (Fig. 2A). Weight loss during R-CDI did not differ between groups (Fig. 2B), but diarrhea onset was delayed in the combination groups: 77% (vancomycin/ramoplanin) and 60% (triple combination) developed diarrhea versus 90% with vancomycin alone (Fig. 2C). Fecal spore shedding was undetectable in all groups on days 5–9 and rose again after treatment ended, with no group differences (Fig. S2C); CFU recovered from intestinal tissue (Fig. 2D–G) and cecal cytotoxicity (Fig. 2H) also did not differ. Together, adding ramoplanin and nystatin to vancomycin delays R-CDI onset.

**Figure 2.**
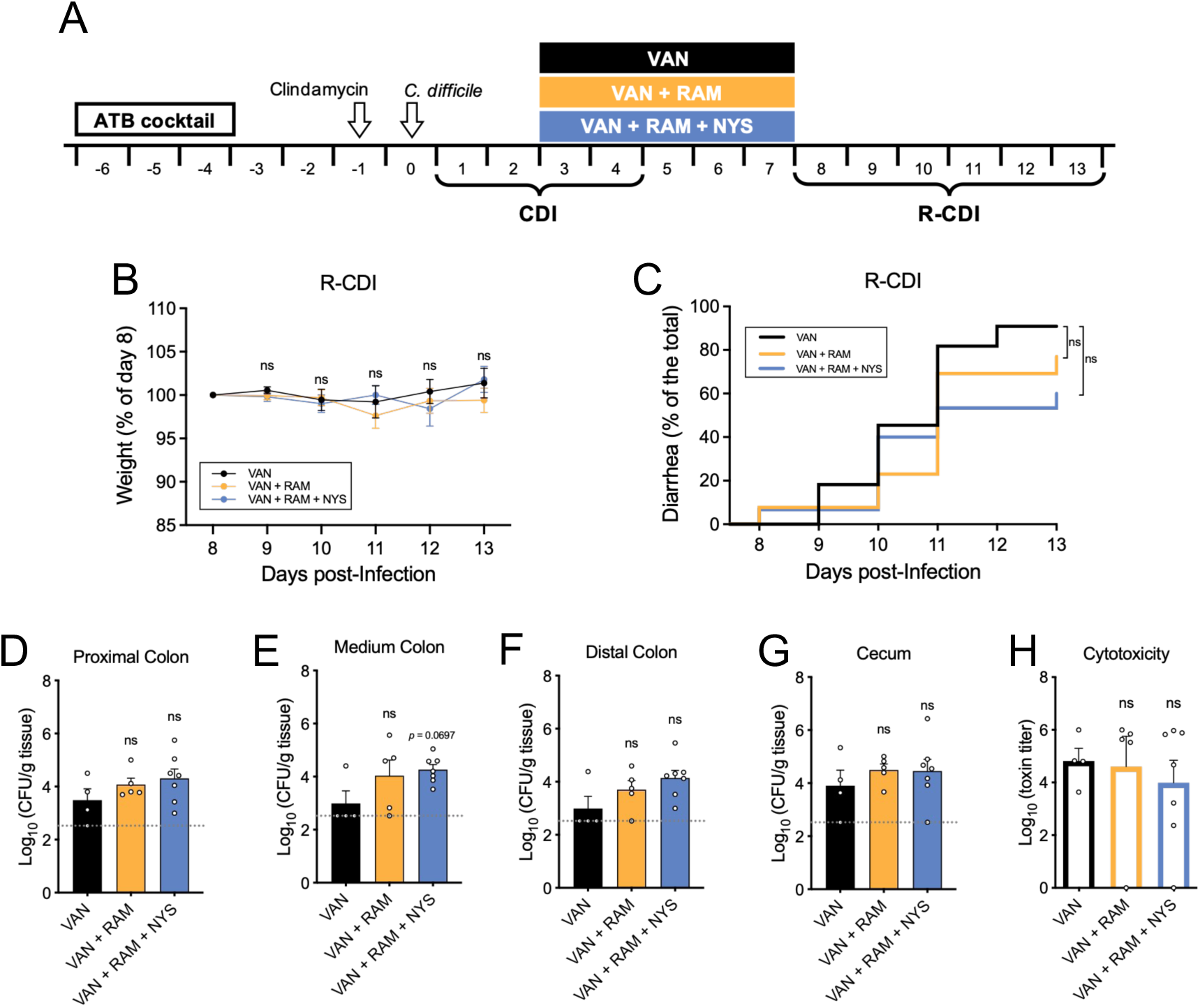

### Induction of germination during antibiotic treatment reduces the incidence of R-CDI

Vancomycin treatment raises intestinal primary bile acids while lowering secondary bile acids ^39^, a shift that favors *C. difficile* spore germination and colonization ^40, 41^. Since germinated spores and vegetative cells are susceptible to antibiotic killing, we reasoned that adding taurocholate to vancomycin and ramoplanin could drive germination of persisting spores for antibiotic clearance, while nystatin limits the reservoir protected from taurocholate. We tested whether adding taurocholate would further delay recurrence. Antibiotic-treated mice were infected with spores (Fig. 3A); both groups had similar weight loss and diarrhea onset during CDI (Fig. S3A, B). At day 3, mice were treated for 5 days with i) vancomycin alone or ii) vancomycin, ramoplanin, nystatin, and taurocholate (Fig. 3A), then monitored for R-CDI. Vancomycin-treated mice showed no weight recovery during R-CDI (Fig. 3B, C), whereas formulation-treated mice regained weight significantly, peaking on day 13 (Fig. 3B, C, D). Only 44% of formulation-treated mice developed R-CDI versus 100% of vancomycin-treated mice (Fig. 3E). Spore shedding (Fig. S3C), intestinal spore load (Fig. 3G–J), and cecal cytotoxicity (Fig. 3K) did not differ. Together, adding taurocholate to the vancomycin/ramoplanin/nystatin formulation reduces the clinical manifestations of R-CDI.

**Figure 3.**
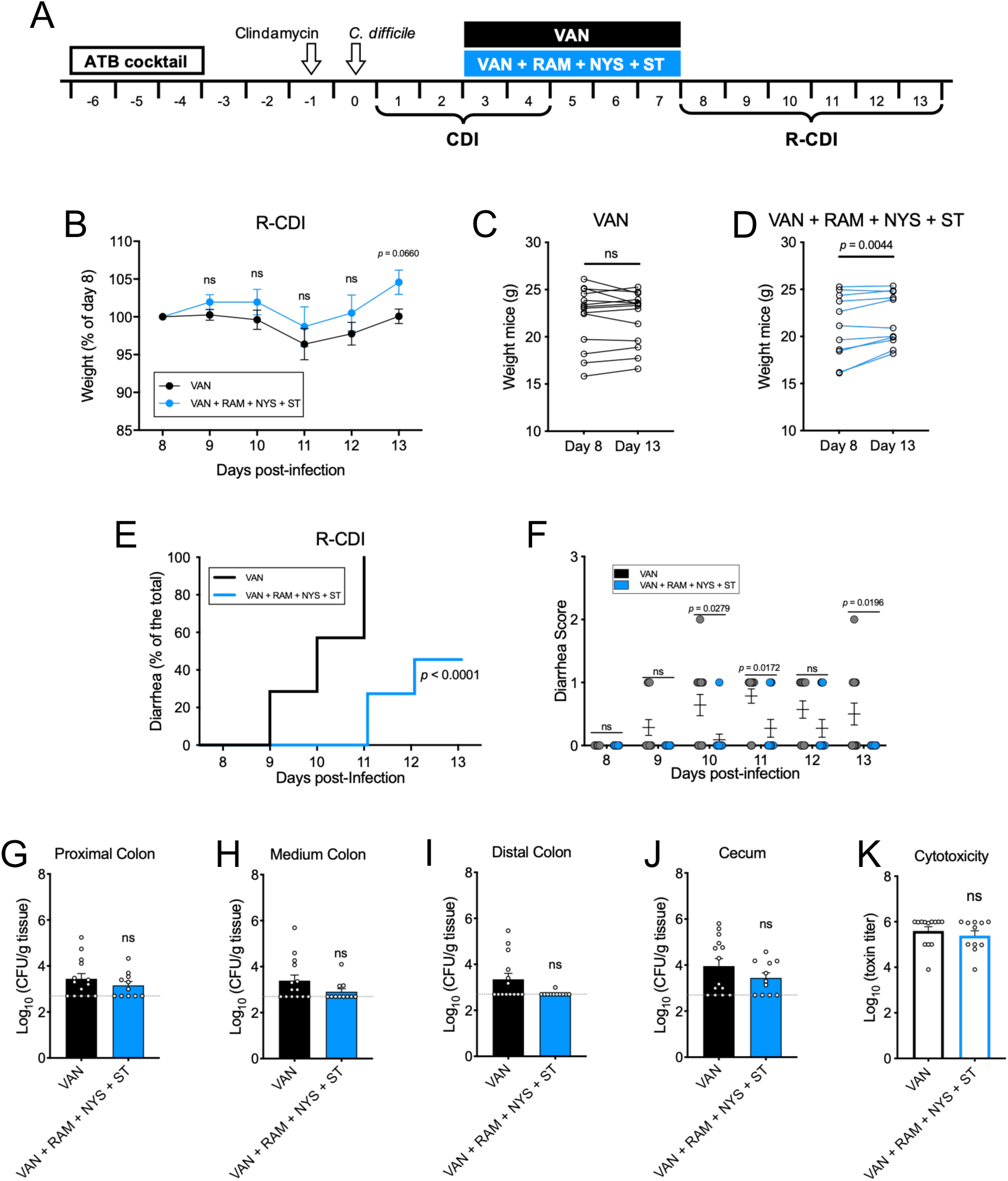

### Vancomycin treatment in combination with ramoplanin, nystatin, and taurocholate reduces the risk of a second R-CDI

Having shown this formulation reduces R-CDI when given during primary CDI, we asked whether giving it during an active R-CDI episode could similarly reduce a second recurrence. Antibiotic-treated mice were infected with spores (Fig. 4A), with similar weight loss and diarrhea onset during CDI (Fig. S4A, B) and during R-CDI development on days 8–13 (Fig. S4C, D); all mice received vancomycin from day 3 for 5 days to induce R-CDI. Mice that developed R-CDI (n = 15) were randomly assigned to two groups and treated on day 13 for 5 days with: i) vancomycin/ramoplanin/nystatin/taurocholate, or ii) vancomycin alone (Fig. 4A), then monitored for a second R-CDI after treatment was suspended. Weight loss during the second R-CDI did not differ between groups (Fig. 4B), but diarrhea onset was significantly delayed with the four-drug formulation (62%) versus vancomycin alone (100%; Fig. 4C). Time to colonization was delayed by one day in the four-drug group, consistent with lower fecal spore load on day 18 (Fig. 4D, E). Notably, mice treated with the four-drug formulation showed higher spore adherence in the medium and distal colon, though not in the proximal colon or cecum content, versus vancomycin alone (Fig. 4F–I); cecal cytotoxicity did not differ (Fig. 4J). Together, the four-drug formulation delays and reduces the clinical manifestations of a second R-CDI.

**Figure 4.**
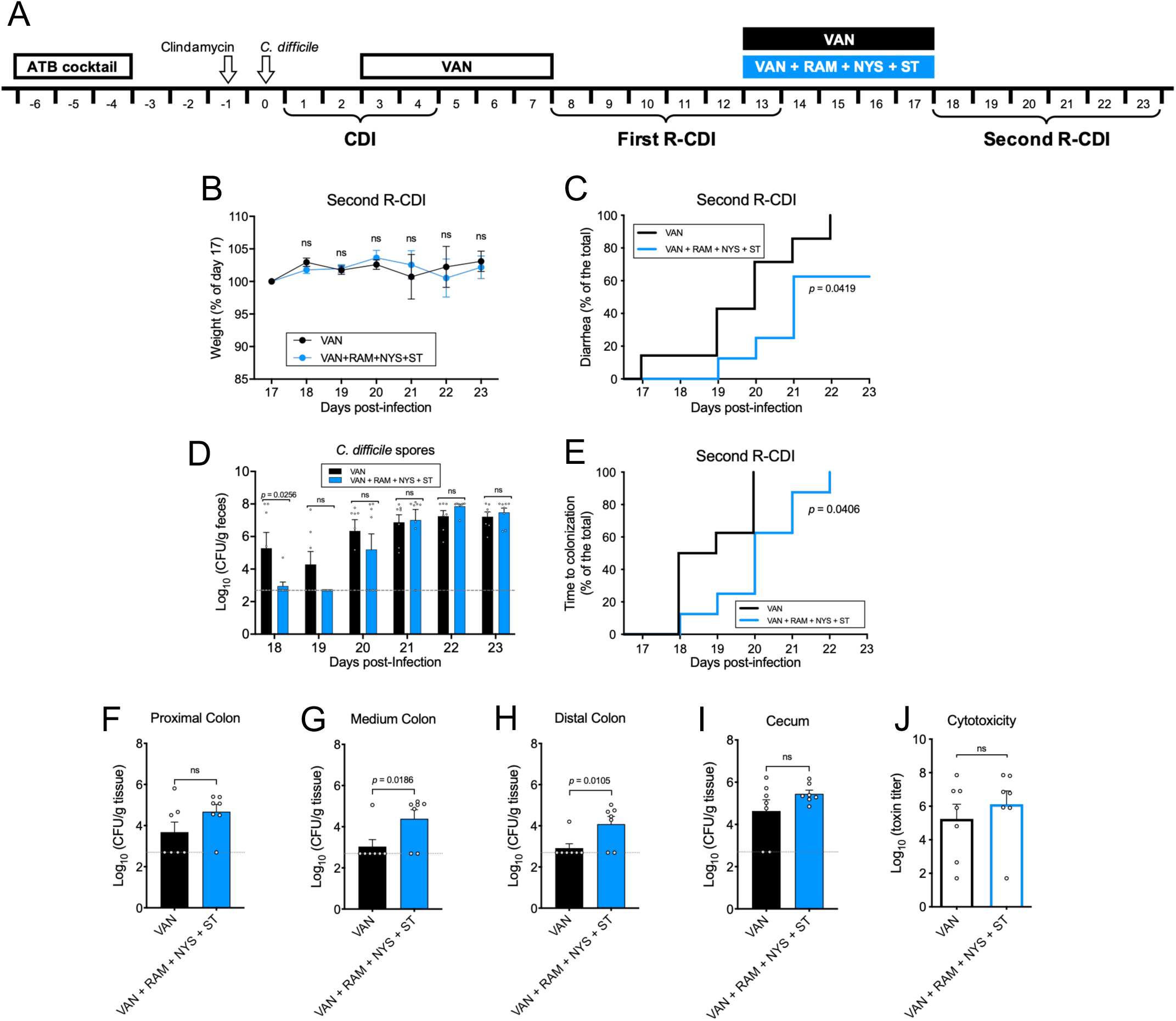

### Vancomycin- and nystatin-loaded chitosan nanoparticles show suitable size, charge, and encapsulation efficiency for mucosal delivery

To increase local intestinal exposure of the combination therapy, we evaluated chitosan nanoparticles containing vancomycin (VAN-CS-NPs) or nystatin (NYS-CS-NPs). Across three independent batches, VAN-CS-NPs averaged 173.4 ± 0.40 nm in diameter with a polydispersity index (PDI) of 0.155 ± 0.013 (Fig. S5A–D), a zeta potential of +16.3 ± 0.2 mV, and a conductivity of 0.0803 mS/cm (Fig. S5E). NYS-CS-NPs averaged 181.3 ± 1.5 nm with a PDI of 0.149 ± 0.003 (Fig. S6A–D), a zeta potential of 15.8 ± 0.03 mV, and a conductivity of 0.0804 mS/cm (Fig. S6E). Encapsulation efficiency by HPLC was 98% for both drugs. Together, both formulations showed a suitable hydrodynamic diameter, positive zeta potential, and homogeneous size distribution.

### Chitosan nanoparticle encapsulation of vancomycin and nystatin outperforms free-drug administration in reducing weight loss, diarrhea, and colonization during R-CDI

Antibiotic-treated mice were infected with C. difficile spores (Fig. 5A), with similar weight loss (Fig. S7A) and time to diarrhea (Fig. S7B) during CDI. On day 3, mice were treated for 5 days with i) vancomycin (n = 9), ii) VAN-CS-NPs (n = 6), iii) vancomycin/nystatin (n = 9), or iv) VAN-CS-NPs/NYS-CS-NPs (n = 10; Fig. 5A), and monitored for R-CDI from day 8. Mice treated with free vancomycin or vancomycin/nystatin showed significant weight loss during R-CDI, peaking on day 11 (Fig. 5B, C, E), whereas mice treated with VAN-CS-NPs or VAN-CS-NPs/NYS-CS-NPs showed no significant weight loss (Fig. 5B, D, F). Maximal weight loss was likewise significantly greater in the free-drug groups (Fig. 5G), consistent with delayed diarrhea onset in nanoparticle-treated groups (Fig. 5H) and reduced diarrhea score on day 10 (Fig. S7C). Spore shedding was reduced on days 10–12 in the nanoparticle-treated groups (Fig. 5I). Using a colonization threshold of 1 × 10 CFU/g of feces, all mice treated with free vancomycin or vancomycin plus nystatin were colonized by day 11 post-infection, compared with significantly lower colonization rates of 33% and 44% in the VAN-CS-NP and VAN-CS-NP plus NYS-CS-NP groups, respectively (Fig. 5J). Spore adherence did not differ between groups in the cecum, proximal colon, or middle colon, but was significantly lower in the distal colon of mice treated with VAN-CS-NPs plus NYS-CS-NPs compared with vancomycin alone (Fig. 5K–N). Cecal cytotoxicity was also significantly lower in both nanoparticle groups versus their respective free-drug groups (Fig. 5O). Together, these results show that nanoparticle-encapsulated vancomycin and nystatin, alone or in combination, are more effective than their free-drug counterparts in reducing weight loss, diarrhea, colonization, and cecal cytotoxicity during R-CDI.

**Figure 5.**
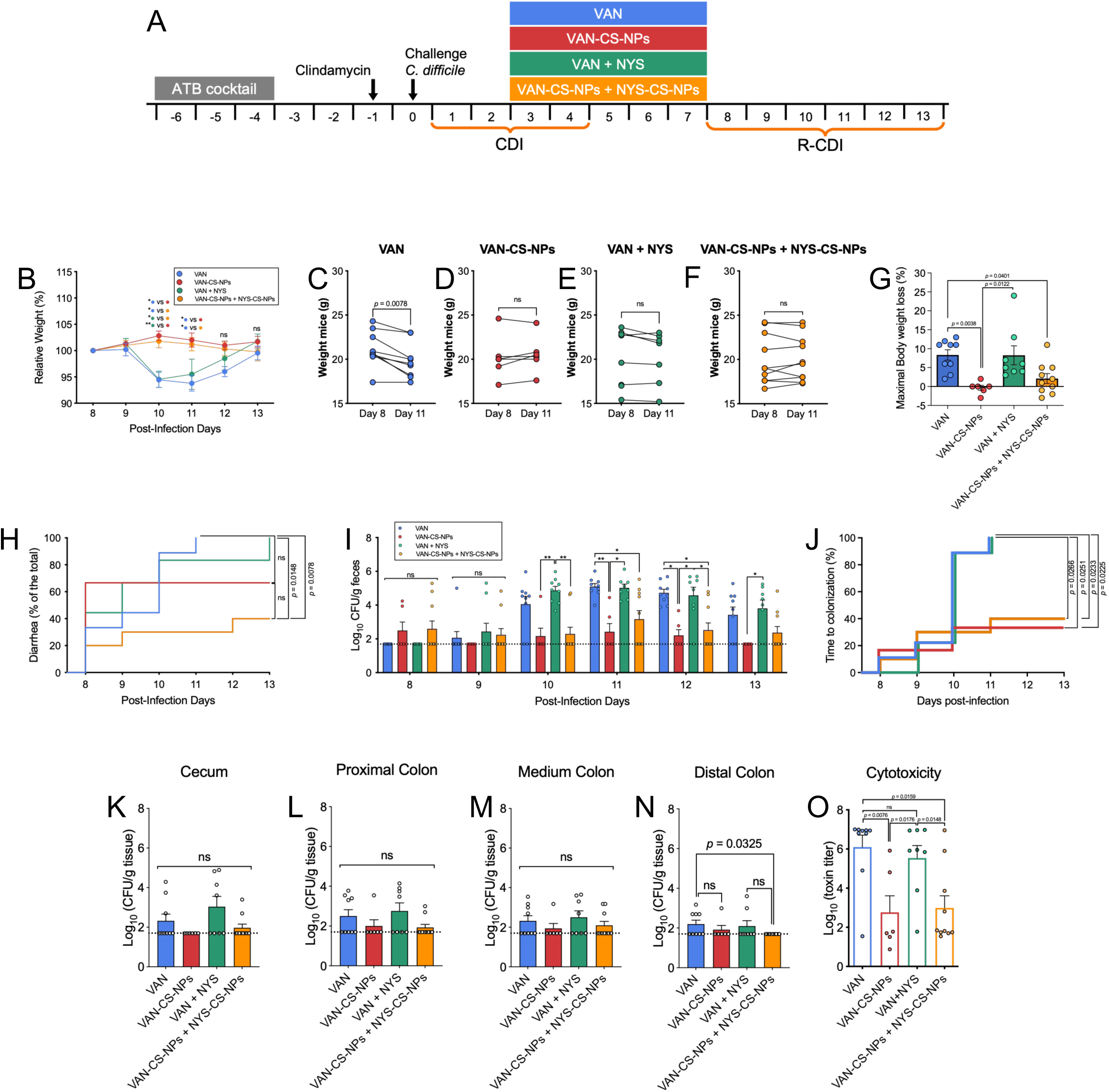

## DISCUSSION

The main finding of this study is that intracellular *C. difficile* spores constitute a therapeutically targetable reservoir contributing to R-CDI. First, we demonstrated that internalized *C. difficile* spores remain metabolically protected from taurocholate-triggered germination, and nystatin reduces their abundance by inhibiting spore internalization. Second, oral administration of a formulation combining vancomycin, ramoplanin, nystatin, and taurocholate during CDI reduces the first recurrence and, when administered during the first recurrence, reduces the second. Third, chitosan nanoparticle delivery of vancomycin and nystatin substantially enhances therapeutic efficacy in animal models relative to free vancomycin.

Internalized *C. difficile* spores remain dormant inside epithelial cells despite extracellular taurocholate and glycine. Extrapolated to an infected patient, this suggests that a subpopulation of spores infecting intestinal epithelial cells may remain dormant during antibiotic treatment of CDI, then be released into the lumen and germinate, contributing to R-CDI. Targeted treatment of this internalized spore reservoir therefore appears crucial to reducing recurrence rates in patients with CDI. Consistent with this, opsonizing spores during CDI has been shown to reduce disease recurrence in preclinical studies ^29^. We recently reported that nystatin inhibits spore internalization into intestinal epithelial cells and the intestinal mucosa ^33^; to our knowledge, no other studies have directly targeted internalized spores as a therapeutic strategy.

Our findings demonstrate that repurposing nystatin, administered alongside antibiotics to mice during CDI, reduces the rate of R-CDI manifestation. Notably, efficiency improved when nystatin was combined only with antibiotics (vancomycin and ramoplanin) to reduce diarrhea during R-CDI.

Adding taurocholate to a formulation of nystatin and antibiotics improved disease resolution by reducing clinical symptoms during R-CDI, though it did not reduce spore adherence to the colonic tract relative to controls. One possible explanation is that intestinal tissue was collected and CFUs assessed 6 days after the formulation was suspended, by which point taurocholate and antibiotic bioavailability had likely declined substantially, permitting colonization. Higher taurocholate concentrations increase *C. difficile* germination rates *in vitro* ^44^, and oral taurocholate has been well tolerated in a small patient cohort at 1 g every 8 h for 2 days ^45^. Increasing colonic taurocholate bioavailability may therefore be important for eliminating germinating spores with antibiotics. Taurocholate is synthesized in the liver by conjugation of unconjugated bile salts (cholate) with taurine; released bile salts are efficiently reabsorbed in the terminal ileum via the apical sodium-dependent bile acid transporter (ASBT) ^46^ and recycled enterohepatically, with only 3-5% lost via feces ^47^. A key limitation of our work is that oral taurocholate may be reabsorbed in the ileum, reducing colonic bioavailability; options to address this include ASBT inhibitors ^46^ or optimizing taurocholate delivery specifically to the colonic mucosa, which is the object of future studies.

We included ramoplanin as a second antibiotic in our formulation because it was among the most promising antibiotics for reducing R-CDI during our experiments; it binds the *C. difficile* spore for a prolonged period, positioning it to attack germinating cells ^28^. In hamster models and preliminary phase II clinical trials, ramoplanin showed effectiveness similar to vancomycin in resolving CDI symptoms ^48, 49^, but to date no phase III trials have been published, so this formulation cannot yet be tested in humans. Future iterations could replace ramoplanin with other antibiotics active against *C. difficile*, such as fidaxomicin, without affecting efficacy.

Nanodelivery of vancomycin and nystatin via chitosan nanoparticles significantly reduced weight loss, diarrhea, colonization, and cytotoxicity relative to free vancomycin or vancomycin plus nystatin. However, whereas we previously found that vancomycin plus free nystatin significantly delayed diarrhea onset from ∼90% to 70% in this model ^33^, the delay observed here when comparing vancomycin alone with vancomycin plus nystatin was non-significant, likely reflecting reduced sample size. Nonetheless, CS-NPs loaded with vancomycin or nystatin represent a promising candidate drug strategy to combat R-CDI. Nanoparticle size, zeta potential, and polydispersity (PDI) shape the nanoparticle-biological interface and in vivo biocompatibility. Nanoparticle size in particular influences pharmacokinetics, tissue distribution, and clearance. Our nanoparticles measured ∼180 nm; particles of 50-100 nm show higher mucus permeability in vitro than 200-nm particles ^50^, suggesting our formulation’s reduced permeability may instead favor increased luminal retention and bioavailability at the site of infection. Chitosan interacts with mucosa electrostatically ^51^, and surface charge (zeta potential) shapes both cytotoxicity and mucoadhesion: cationic particles are generally more cytotoxic, though this also depends on cell and membrane type, while neutral or positively charged ∼100-nm particles show greater mucus permeability and retention than larger or negatively charged particles ^52^, and zeta potential and size are inversely related ^53^. Our particles’ positive zeta potential (+15.8 ± 0.03 mV) likely favors electrostatic interaction between chitosan’s positive charges and the negatively charged microbial membrane ^54, 55^, consistent with reports that particle size and zeta potential are key determinants of nanoparticle antimicrobial activity. Our PDI of ∼0.15 falls within the acceptable range (monodisperse below 0.05; broadly polydisperse above 0.7) ^52^.

In conclusion, this study establishes a multifactorial therapeutic framework combining inhibition of spore internalization, induction of dormant-spore germination, bacterial killing, and enhanced mucoadhesive nanoparticle delivery. This approach substantially reduces R-CDI burden in preclinical models and offers a cost-effective, orally administered alternative for reducing recurrence. Future work should optimize taurocholate delivery, explore alternative antibiotics, and evaluate this formulation in clinical trials to establish safety and efficacy in humans.

## Notes

The authors declare no competing interests.

## Supplementary information

The following supporting information is available free of charge at ACS website http://pubs.acs.org.

Figure S1. Effect of heat treatment on the germination of *C. difficile* spores in monolayers of Caco-2 cells.

Figure S2. Supplementary figures for the R-CDI experiment detailed in Figure 2.

Figure S3. Supplementary figures for the R-CDI experiment details in Figure 3.

Figure S4. Supplementary figures for the second R-CDI experiment are shown in Figure 4.

Figure S5. Physicochemical characterization of VAN-CS-NPs.

Figure S6. Physicochemical characterization of NYS-CS-NPs.

Figure S7. Supplementary data related to Fig. 5.

## Authoŕs Contribution

**<u>Conceptual Design</u>** (Formulation of hypothesis, development of study objectives, defining experimental, statistical and analytical procedures:

**DPS, PCC, CO, FG**

**<u>Data acquisition</u>** (labwork, theoretical calculations, literature searches)

**PCC, MML, NMB, JMM, FDY, FL, DC, RH-A, JV, KM,**

**<u>Analysis and interpretation of the data</u>** (making sense of and presenting the results, data analysis (statistical or other)

**PCC, MML, NMB, JMM, FDY, FL, DC, RH-A, JV, KM, DPC, CO, FG**

**<u>Writing publication</u>** (creating all or substantial part of the scientific and technical product)

**PCC, CO, FG, MPG, DPS**

**<u>Critical Revision of Publication</u>** (reworking all the scientific and technical product for intellectual content before submission (not just spelling and grammar checking)

**PCC, CO, FG, MPG, DPS**

**<u>Approval of final publication</u>** (providing approval of the final product to be published)

**DPS**

**<u>Supervision</u>** (oversight and responsibility for the study, general supervision of the research group)

**DPS, CO**

**<u>Resources</u>** (funding, equipment, facilities, personnel vital to the project)

**DPS**

## REFERENCES

1. Gerding DN, Johnson S, Peterson LR et al. *Clostridium difficile*-associated diarrhea and colitis. Infect Control Hosp Epidemiol 1995; 16: 459–77.

2. Nasiri MJ, Goudarzi M, Hajikhani B et al. *Clostridioides* (*Clostridium*) *difficile* infection in hospitalized patients with antibiotic-associated diarrhea: A systematic review and meta-analysis. Anaerobe 2018; 50: 32–7.

3. Evans CT, Safdar N. Current Trends in the Epidemiology and Outcomes of *Clostridium difficile* Infection. Clin Infect Dis 2015; 60 **Suppl 2**: S66–71.

4. Centers for Disease Control and Prevention. Antibiotic Resistance Threats in the United States, 2019. Accessed Dec 19, 2025. https://www.cdc.gov/antimicrobial-resistance/media/pdfs/2019-ar-threats-report-508.pdf.

5. McFarland LV, Elmer GW, Surawicz CM. Breaking the cycle: treatment strategies for 163 cases of recurrent *Clostridium difficile* disease. Am J Gastroenterol 2002; 97: 1769–75.

6. McDonald LC, Gerding DN, Johnson S et al. Clinical Practice Guidelines for *Clostridium difficile* Infection in Adults and Children: 2017 Update by the Infectious Diseases Society of America (IDSA) and Society for Healthcare Epidemiology of America (SHEA). Clin Infect Dis 2018; 66: 987–94.

7. Bouza E. Consequences of *Clostridium difficile* infection: understanding the healthcare burden. Clin Microbiol Infect 2012; 18 **Suppl 6**: 5–12.

8. Olsen MA, Yan Y, Reske KA et al. Recurrent *Clostridium difficile* infection is associated with increased mortality. Clin Microbiol Infect 2015; 21: 164–70.

9. Seiffert A, Noack S, Wingen-Heimann S et al. Epidemiology and Course of Clostridioides difficile Infections in Germany: Retrospective Analysis of German Health Claims Data. Infect Dis Ther 2025; 14: 2341–57.

10. Kelly CP. Can we identify patients at high risk of recurrent *Clostridium difficile* infection? Clin Microbiol Infect 2012; 18 **Suppl 6**: 21–7.

11. Giau VV, Lee H, An SSA et al. Recent advances in the treatment of *C. difficile* using biotherapeutic agents. Infect Drug Resist 2019; 12: 1597–615.

12. Schaffler H, Breitruck A. *Clostridium difficile* - From Colonization to Infection. Front Microbiol 2018; 9: 646.

13. Ducarmon QR, Zwittink RD, Hornung BVH et al. Gut Microbiota and Colonization Resistance against Bacterial Enteric Infection. Microbiol Mol Biol Rev 2019; 83.

14. Lyras D, O’Connor JR, Howarth PM et al. Toxin B is essential for virulence of *Clostridium difficile*. Nature 2009; 458: 1176–9.

15. Kuehne SA, Cartman ST, Heap JT et al. The role of toxin A and toxin B in *Clostridium difficile* infection. Nature 2010; 467: 711–3.

16. Paredes-Sabja D, Shen A, Sorg JA. *Clostridium difficile* spore biology: sporulation, germination, and spore structural proteins. Trends Microbiol 2014; 22: 406–16.

17. Deakin LJ, Clare S, Fagan RP et al. The *Clostridium difficile spo0A* gene is a persistence and transmission factor. Infect Immun 2012; 80: 2704–11.

18. Sethi AK, Al-Nassir WN, Nerandzic MM et al. Persistence of skin contamination and environmental shedding of *Clostridium difficile* during and after treatment of *C. difficile* infection. Infect Control Hosp Epidemiol 2010; 31: 21–7.

19. Polivkova S, Krutova M, Capek V et al. Fidaxomicin versus metronidazole, vancomycin and their combination for initial episode, first recurrence and severe *Clostridioides difficile* infection - An observational cohort study. Int J Infect Dis 2021; 103: 226–33.

20. Stevens VW, Nelson RE, Schwab-Daugherty EM et al. Comparative Effectiveness of Vancomycin and Metronidazole for the Prevention of Recurrence and Death in Patients With *Clostridium difficile* Infection. JAMA Intern Med 2017; 177: 546–53.

21. Cornely OA, Nathwani D, Ivanescu C et al. Clinical efficacy of fidaxomicin compared with vancomycin and metronidazole in *Clostridium difficile* infections: a meta-analysis and indirect treatment comparison. J Antimicrob Chemother 2014; 69: 2892–900.

22. Al Momani LA, Abughanimeh O, Boonpheng B et al. Fidaxomicin vs Vancomycin for the Treatment of a First Episode of *Clostridium difficile* Infection: A Meta-analysis and Systematic Review. Cureus 2018; 10: e2778.

23. Feuerstadt P, Louie TJ, Lashner B et al. SER-109, an Oral Microbiome Therapy for Recurrent Clostridioides difficile Infection. N Engl J Med 2022; 386: 220–9.

24. Villano SA, Seiberling M, Tatarowicz W et al. Evaluation of an oral suspension of VP20621, spores of nontoxigenic *Clostridium difficile* strain M3, in healthy subjects. Antimicrob Agents Chemother 2012; 56: 5224–9.

25. Gerding DN, Meyer T, Lee C et al. Administration of spores of nontoxigenic Clostridium difficile strain M3 for prevention of recurrent C. difficile infection: a randomized clinical trial. JAMA 2015; 313: 1719–27.

26. Somner EA, Reynolds PE. Inhibition of peptidoglycan biosynthesis by ramoplanin. Antimicrob Agents Chemother 1990; 34: 413–9.

27. Lo M-C, Men H, Branstrom A et al. A new mechanism of action proposed for ramoplanin. Journal of the American Chemical Society 2000; 122: 3540–1 %@ 0002-7863.

28. Kraus CN, Lyerly MW, Carman RJ. Ambush of *Clostridium difficile* spores by ramoplanin: activity in an in vitro model. Antimicrob Agents Chemother 2015; 59: 2525–30.

29. Pizarro-Guajardo M, Díaz-González F, Álvarez-Lobos M et al. Characterization of Chicken IgY Specific to *Clostridium difficile* R20291 Spores and the Effect of Oral Administration in Mouse Models of Initiation and Recurrent Disease. Front Cell Infect Microbiol 2017; 7: 365.

30. www.drugs.com. Vowst Prices, Coupons, Copay Cards & Patient Assistance. https://www.drugs.com/price-guide/vowst.

31. Wilcox MH, Gerding DN, Poxton IR et al. Bezlotoxumab for Prevention of Recurrent *Clostridium difficile* Infection. N Engl J Med 2017; 376: 305–17.

32. Fan AE, Alqahtani S, Garnes ND et al. Bezlotoxumab as Treatment for Recurrent/Chronic Clostridioides difficile Infection in Pediatric Stem Cell Transplant Recipients: A Multi-Institutional Experience. Pediatr Blood Cancer 2025; 72: e31896.

33. Castro-Córdova P, Mora-Uribe P, Reyes-Ramírez R et al. Entry of spores into intestinal epithelial cells contributes to recurrence of *Clostridioides difficile* infection. Nat Commun 2021; 12: 1140.

34. U.S. Food and Drug Administration. Application letter and printed labeling for nystatin. https://www.accessdata.fda.gov/drugsatfda_docs/anda/98/64142ap_appltr_prntlbl_chemr.pdf.

35. Calderon-Romero P, Castro-Cordova P, Reyes-Ramirez R et al. *Clostridium difficile* exosporium cysteine-rich proteins are essential for the morphogenesis of the exosporium layer, spore resistance, and affect *C. difficile* pathogenesis. PLoS Pathog 2018; 14: e1007199.

36. Chen X, Katchar K, Goldsmith JD et al. A mouse model of *Clostridium difficile*-associated disease. Gastroenterology 2008; 135: 1984–92.

37. Castro-Córdova P, Díaz-Yáñez F, Muñoz-Miralles J et al. Effect of antibiotic to induce *Clostridioides difficile*-susceptibility and infectious strain in a mouse model of *Clostridioides difficile* infection and recurrence. Anaerobe 2020; 62: 102149.

38. Warren CA, van Opstal EJ, Riggins MS et al. Vancomycin treatment’s association with delayed intestinal tissue injury, clostridial overgrowth, and recurrence of *Clostridium difficile* infection in mice. Antimicrob Agents Chemother 2013; 57: 689–96.

39. Vrieze A, Out C, Fuentes S et al. Impact of oral vancomycin on gut microbiota, bile acid metabolism, and insulin sensitivity. J Hepatol 2014; 60: 824–31.

40. Theriot CM, Bowman AA, Young VB. Antibiotic-Induced Alterations of the Gut Microbiota Alter Secondary Bile Acid Production and Allow for *Clostridium difficile* Spore Germination and Outgrowth in the Large Intestine. mSphere 2016; 1.

41. Theriot CM, Koenigsknecht MJ, Carlson PE et al. Antibiotic-induced shifts in the mouse gut microbiome and metabolome increase susceptibility to *Clostridium difficile* infection. Nat Commun 2014; 5: 3114.

42. Sorg JA, Sonenshein AL. Bile salts and glycine as cogerminants for *Clostridium difficile* spores. J Bacteriol 2008; 190: 2505–12.

43. Walters BA, Roberts R, Stafford R et al. Relapse of antibiotic associated colitis: endogenous persistence of *Clostridium difficile* during vancomycin therapy. Gut 1983; 24: 206–12.

44. Sorg JA, Sonenshein AL. Inhibiting the initiation of *Clostridium difficile* spore germination using analogs of chenodeoxycholic acid, a bile acid. J Bacteriol 2010; 192: 4983–90.

45. Plusa SM, Clark NW. Prevention of postoperative renal dysfunction in patients with obstructive jaundice: a comparison of mannitol-induced diuresis and oral sodium taurocholate. J R Coll Surg Edinb 1991; 36: 303–5.

46. Slijepcevic D, van de Graaf SF. Bile Acid Uptake Transporters as Targets for Therapy. Dig Dis 2017; 35: 251–8.

47. Dawson PA, Karpen SJ. Intestinal transport and metabolism of bile acids. J Lipid Res 2015; 56: 1085–99.

48. Freeman J, Baines SD, Jabes D et al. Comparison of the efficacy of ramoplanin and vancomycin in both in vitro and in vivo models of clindamycin-induced *Clostridium difficile* infection. J Antimicrob Chemother 2005; 56: 717–25.

49. Gerding DN, Johnson S. Management of *Clostridium difficile* infection: thinking inside and outside the box. Clin Infect Dis 2010; 51: 1306–13.

50. Bandi SP, Kumbhar YS, Venuganti VVK. Effect of particle size and surface charge of nanoparticles in penetration through intestinal mucus barrier.

51. He Ping, Davis Stanley S., Lisbeth I. In vitro evaluation of the mucoadhesive properties of chitosan microspheres. International Journal of Pharmaceutics 1998; 75–88.

52. Lai SK, Wang YY, Hanes J. Mucus-penetrating nanoparticles for drug and gene delivery to mucosal tissues. Adv Drug Deliv Rev 2009; 61: 158–71.

53. Syed Wazed ALI, Mangala J, Subbiyan R. S ynthesis and characterization of chitosan nanoparticles with enhanced antimicrobial activity. International Journal of Nanoscience 2011; 10: 979–84.

54. Qi L, Xu Z, Jiang X et al. Preparation and antibacterial activity of chitosan nanoparticles. Carbohydr Res 2004; 339: 2693–700.

55. Ing LY, Zin NM, Sarwar A et al. Antifungal activity of chitosan nanoparticles and correlation with their physical properties. Int J Biomater 2012; 2012: 632698.

